# Novitski’s Distal shift in Paracentric Inversion Evolution

**DOI:** 10.1101/485334

**Authors:** Spencer A. Koury

## Abstract

In *Drosophila pseudoobscura* younger chromosomal inversions tend to be found distal to older inversions. By examining phylogenetic series of overlapping inversions for 134 gene arrangements of 13 chromosomes this pattern was extended to five additional *Drosophila* species. This distinct pattern arose repeatedly and independently in all six species and likely reflects an underlying principle of chromosome evolution. In this study it is illustrated how transmission of distal inversions is *always* favored in female meiosis when crossing over in homosequential regions of overlapping inversions generates asymmetric dyads. This cytogenetic mechanism for female meiotic drive is described in detail and advanced as an explanation for the distal shift in phylogenetic series of overlapping inversions as well as several better known patterns in the evolution of serially inverted chromosomes.

## INTRODUCTION

The study of *Drosophila* chromosomal inversion polymorphism emerged as model system for evolutionary genetics in the 1930’s and played a major role in the Modern Synthesis (Krimbas and Powell, 1992). Interest in inversions started to wane in the 1960’s with the appearance of genetic markers, such as allozymes, that could be more readily applied to many other organisms. However, beginning in the 1990’s, application of sequencing technology in natural population genetics led to the discovery of inversions in several non-model organisms and a renaissance in both empirical and theoretical research on inversion polymorphism and its role in adaptation and speciation (Jones *et al.*, 2012; Kirkpatrick and Barton, 2006; Le Poul *et al.*, 2014; Lowry and Willis, 2010). Clearly, inversion polymorphism is an aspect of structural genome evolution in more than just *Drosophila*. However, for the investigation of nuanced patterns and mechanisms in chromosome evolution the knowledge base developed in *Drosophila* cytogenetics remains indispensable (Corbett-Detig and Hartl, 2012; Schaeffer *et al.*, 2008). As information on inversion polymorphism accumulates in the post-genomic era, early studies of chromosomal rearrangements in model systems will play a fundamental role in organizing these abundant data to uncover the principles of chromosome evolution (Bhutkar *et al.*, 2008; Gong *et al.*, 2005; Ranz *et al.*, 2007).

Inversion polymorphism in *Drosophila* was the first genetic marker system studied for phylogenetic inference and natural population genetics. Central to this research was Alfred Sturtevant, who first discovered chromosomal inversions, their effects in transmission genetics, and described natural variation in several species (Sturtevant and Novitski, 1941; Sturtevant, 1917; Sturtevant, 1921; Sturtevant and Beadle, 1936; Sturtevant and Dobzhansky, 1936). In 1936, Sturtevant proposed to make inversion polymorphism in *D. pseudoobscura* a model system for evolutionary biology through collaboration with Theodosius Dobzhansky and Sewall Wright (Provine, 1981). From this collaboration came the first phylogenetic series of overlapping inversions (figure 1), and would for the next four decades be expanded by Dobzhansky *et alia* into the Genetics of Natural Populations I-XLIII (Dobzhansky and Sturtevant, 1938; Lewontin, 1981).

**Figure 1.**
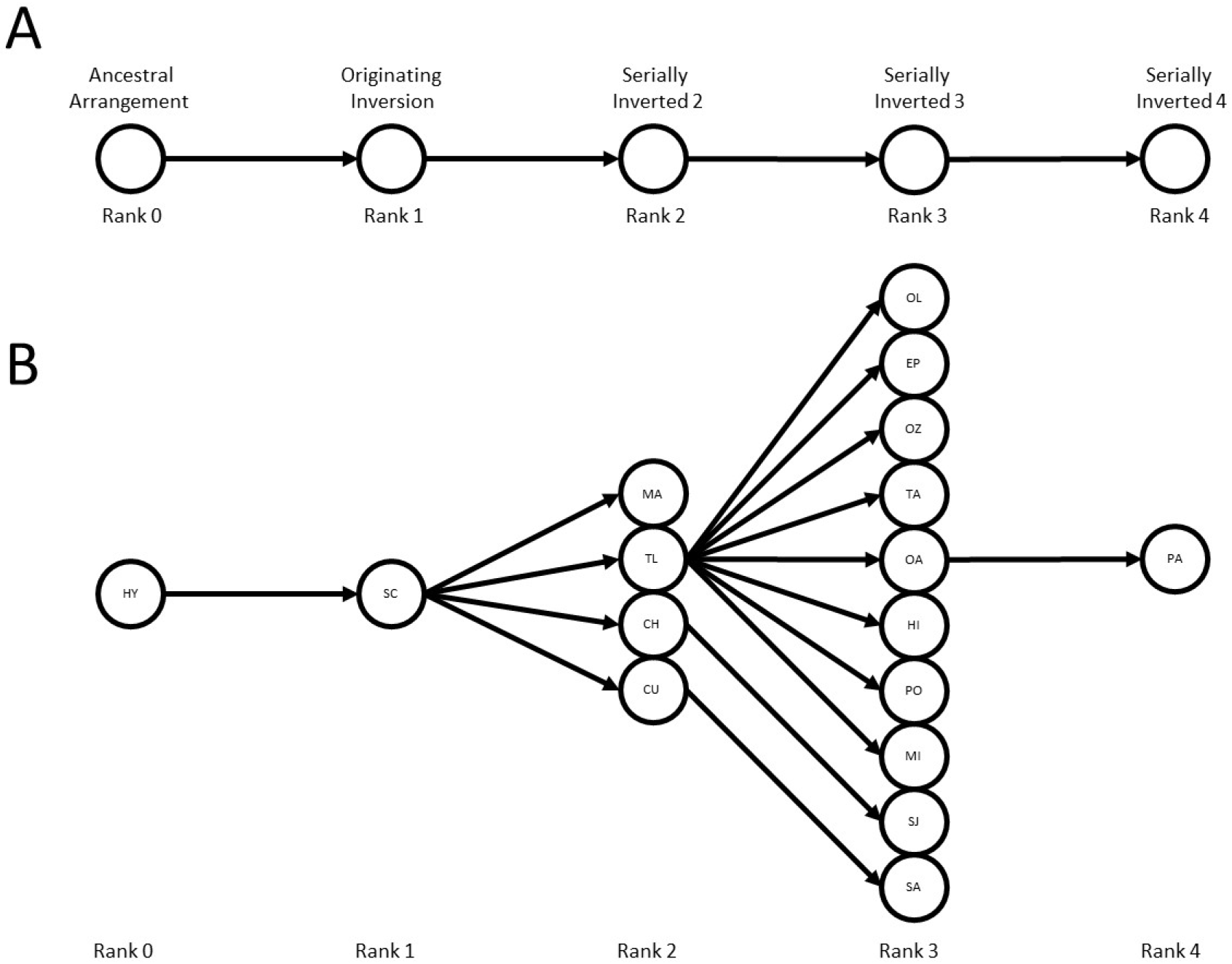
Schematics of phylogenetic series. A) A simple series from ancestral arrangement (rank 0), to the series originating inversion (rank 1), and then serially inverted chromosomes of rank 2 through 4. B) The “Santa Cruz” phylogenetic series of *D. pseudoobscura* chromosome III (illustrating only arrangements repeatedly observed in natural populations). Two letter codes refer to names of each gene arrangement (*cf.* Dobzhansky and Epling, 1944).

Despite this impressive scientific history, the mechanisms governing inversion origin, establishment, and maintenance remain obscure. The experimental study of spontaneous mutation of inversions and their invasion in populations is logistically difficult, if not impossible, and thus remains underexplored (Krimbas and Powell, 1992; Yamaguchi and Mukai, 1974). Balancing selection (associative overdominance, multiple niche, *etc*.) has long been favored in the maintenance of inversion polymorphism and has been variously supported by sampling natural populations as well as population cage and field experiments (Dobzhansky, 1948; Levitan and Etges, 2009; Schaeffer, 2008; Wright and Dobzhansky, 1946). However, experimental efforts on all fronts are intrinsically biased by the idiosyncrasies of chromosomal breakages and linked genetic backgrounds for the small handful of common inversions that provide the genetic material for analysis (*e.g.* Dobzhansky, 1950; Levitan, 1962; Yamaguchi and Mukai, 1974).

In contrast, non-experimental methods such as direct sequencing of inversion breakpoints and surveys of molecular diversity in inverted regions have provided a historical perspective on the origin of the common inversions. Common cosmopolitan inversions of *D. melanogaster* have reduced levels of polymorphism and originated relatively recently, on the order of 100,000 years ago (Andolfatto *et al.*, 1999; Corbett-Detig and Hartl, 2012; Hasson and Eanes, 1996; Matzkin *et al.*, 2005; Wesley and Eanes, 1994). Similar patterns are observed for the serially inverted third chromosome of *D. pseudoobscura*, where the long and complex history of local adaptation and selection of epistatic effects have shaped the molecular diversity among gene arrangements (Fuller *et al.*, 2016; Schaeffer, 2008; Schaeffer *et al.*, 2003; Wallace *et al.*, 2011; Wallace *et al.*, 2013). However, the discovery that common inversions from several species exhibit long range linkage disequilibrium, epistatic fitness effects, and are associated with meiotic drive renews concerns about drawing inferences from the exclusive study of this relatively small sample of inversion polymorphism (Corbett-Detig and Hartl, 2012; Houle and Márquez, 2015; Schaeffer *et al.*, 2003).

A long standing hypothesis that inversions result from ectopic recombination of transposable elements or other repetitive sequence found little support in the first polymorphic inversion breakpoints to be directly sequenced, *D. melanogaster*’s *In(3L)P*, *In(2L)t*, and *In(3R)P* (Andolfatto *et al.*, 1999; Matzkin *et al.*, 2005; Wesley and Eanes, 1994). Outside of *D. melanogaster*, there is both direct (Cáceres *et al.*, 1999b) and indirect (Orengo *et al.*, 2015) support of this mechanism. However, in 29 fixed inversions in the *melanogaster* group, Ranz *et al.* (2007) found only two instances of inverted repetitive sequences that would even allow this mechanism to operate. Complicating this historical analysis is the tendency for transposable elements to accumulate on the minority arrangements (Eanes *et al.*, 1992; Sniegowski and Charlesworth, 1994), and the possibility of transposable element remnants eroding over time (Puerma *et al.*, 2014; Ranz *et al.*, 2007). No unified characterization of inversion breakpoints, or the mechanisms governing the process, has emerged as the breakpoints studied have ranged from simple “cut and paste” to complex rearrangements including small inverted duplications and deletions (Cáceres *et al.*, 1999a; Ranz *et al.*, 2007; Wesley and Eanes, 1994).

Beyond the molecular characterization, the cytological study of inversion breakpoints along chromosome arms can also provide information about mechanisms controlling inversion polymorphism. The standard null distribution would assume inversions result from a rejoining of two chromosomal breakages that occur with uniform probability along the chromosome (Van Valen and Levins, 1968). When compared to this null distribution, breakpoints of a given inversion are observed to be further apart than expected, creating a deficiency of small inversions (*e.g.* Brehm and Krimbas, 1991). When comparing breakpoints *among* inversions, the breakpoints tend to be grouped closer together than expected creating extensive overlap of chromosomal inversions (Novitski, 1946). Finally, rather than the expected uniform distribution, serially inverted chromosomes tend to have inversion breakpoints clustered in distal regions of chromosome arms (Novitski, 1946).

Using the direct ancestor-descendant relationships of gene arrangements within species to generate phylogenetic series, it can be shown that breakpoints of derived inversions tend to lie distal of the corresponding inversion breakpoints in the ancestral arrangement (Novitski, 1946; Sturtevant and Dobzhansky, 1936). The telomeric progression of overlapping rearrangements for serially inverted chromosomes is called Novitiski’s distal shift, because this little-known rule of chromosome evolution was first described by Ed Novitski with cytogenetic evidence from Muller element C of *Drosophila pseudoobscura* (Krimbas and Powell, 1992; Novitski, 1946). o extend the observation of Novitski’s distal shift, I examined published data from 28 phylogenetic series for 13 different chromosomes in six *Drosophila* species of the *obscura* group. Each phylogenetic series arose independently in each species and represents the direct ancestor-descendent relationship in paracentric inversion evolution. Having validated the distal shift empirically, I propose a meiotic drive mechanism for the evolution of overlapping chromosomal inversions that explains the progressive distal shift towards the telomere. I discuss difficulties in extending this mechanism to serially inverted chromosomes and suggest the same mechanism is the cause of other notable patterns in paracentric inversion evolution.

## MATERIALS AND METHODS

The cytogenetics for sixteen *Drosophila* species in the *obscura* group have been published. Although most species in this group have extensive inversion polymorphism, only six species had sufficient data (multiple overlapping rearrangements) to allow the construction of phylogenies for overlapping paracentric inversions. Inversion phylogenies are unrooted trees. Incorporating information on arrangement frequency, geographical distribution, and the karyotype of interspecific hybrids, consensus ancestral arrangements (often designated “Standard”) are used to polarize inversion phylogenies. Note, arrangements other than the Standard may have been ancestral, as proposed for two chromosomes in *D. subobscura* and demonstrated with molecular variation for *D. pseudoobscura* (Krimbas, 1992; Wallace *et al.*, 2011). The first gene rearrangement in a phylogenetic series is designated as the “series originating inversion.” An inversion’s rank in a phylogenetic series was recorded as the minimal number of inversion back steps required to obtain the ancestral arrangement. Species and chromosome information for the 28 phylogenetic series of overlapping inversions used in this paper are listed in table 1 with a sample series illustrated in figure 1 (see supplementary figure 1 for dataset from all series)

**Table 1.**
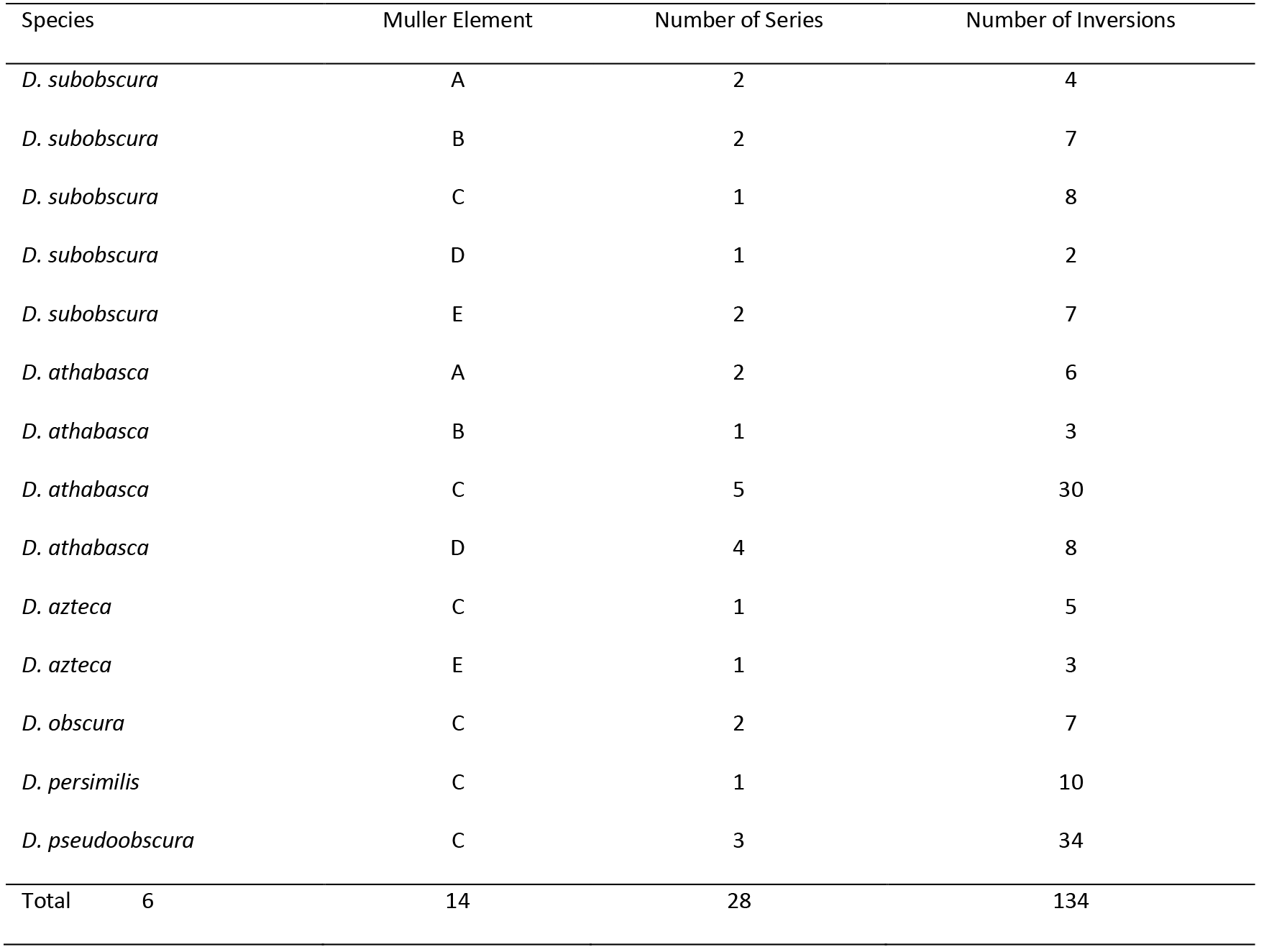
The list of 28 phylogenetic series used in this study. Note that the 14 different Muller elements listed constitute only 13 chromosomes as elements A and D were fused to form the metacentric X chromosome in *D. athabasca.*

To quantify the distal shift, the cytogenetic location of breakpoints were converted to a numerical value where the centromere position was zero and each successive cytogenetically discernable region is considered one unit distal. Where ambiguity exists for an inversion breakpoint location, the midpoint of the range indicated was taken. To facilitate comparison across chromosomes and species, the scale for each chromosome was standardized to 100 unit lengths. This quantification does not necessarily correspond to a linear function of physical distance (bp) or genetic distance (cM), but it does assign a value between 0-100 for every inversion breakpoint and represents its cytogenetic location on the chromosome relative to the centromere. Inversion breakpoints location on this scale must be determined based on the arrangement upon which that new inversion first arose, not necessarily on the standard arrangement scale that is often reported in the literature. Similarly, inversion size was estimated as the distance between proximal and distal inversion breakpoints on the gene arrangement upon which it first occurred. Thus breakpoint location, inversion size, and phylogenetic rank for a given inversion are not obvious from casual observation and these data are provided in supplemental table 1.

To test the distal shift, inversion breakpoint location and size must be adjusted for the position and size of each respective phylogenetic series. Inversion breakpoint locations were therefore expressed as a deviation from the midpoint of the series originating inversion and inversion size was expressed as a deviation from the average inversion size for each phylogenetic series. On the standardized adjusted scale, zero represented the location of the originating inversion for each series, positive values represented a distal shift, and negative values indicated proximal movement of the derived inversion. Similarly, positive values indicates an above average size and negative values represent a smaller than average size after standardization and adjustment for each phylogenetic series.

Statistical analysis of the distal shift was performed by least squares regression of standardized inversion breakpoint location upon the inversion’s rank in a phylogenetic series. For each phylogenetic series the Muller element and species were incorporated as nested categorical variables in the regression analysis. Phylogenetic correction for the species term (incorporating non-independence of observations due to shared ancestry) is statistically inappropriate when analyzing polymorphisms unique to each species, precisely because there can be no covariation due to coancestry. Separate analyses were performed for proximal and distal inversion breakpoints with statistical significance of the regression coefficients assessed by two sided *t*-test for *β = 0*. A statistical test for equality of slopes was performed using *F*-statistics for the null hypothesis *β_proximal_* = *β_distal_* (Sokal and Rohlf, 1995). Statistical significance of inversion size reduction was assessed by two sided *t*-test for *β = 0* after regression of standardized inversion size on phylogenetic series.

## RESULTS

The quantitative analysis of 134 inversions in 28 phylogenetic series on 13 chromosomes from six *obscura* group species provided strong statistical support of the telomeric progression of sequentially derived inversion breakpoints (figure 2). The regression of proximal inversion breakpoint location on phylogenetic rank yielded a statistically significant regression coefficient (*β=10.00, t=4.54, df=1, p< 0.001*) (table 2). Distal inversion breakpoint location when regressed on phylogenetic rank was also statistically significant (*β=7.07, t=2.98, df=1 p=0.004*) (table 3). Variance components and the occasional statistical significance associated with nested categorical variables (species, element, series) is likely a product of standardization across chromosomes with heterogeneous map resolution and genetic length. Again phylogenetic correction for these tests is both logically and statistically inappropriate as the inversion polymorphism analyzed is unique to and arose independently in each species.

**Table 2.** ANOVA table for regression of proximal inversion breakpoint location on phylogenetic rank. Data corresponds to the open symbols in figure 2.

| Source of Variation | df | SS | MS | F ratio | p |
| --- | --- | --- | --- | --- | --- |
| Regression on Rank | 1 | 5149.37 | 5149.37 | 20.58 | < 0.001 |
| Species | 5 | 2806.09 | 561.22 | 2.24 | 0.055 |
| Element within Species | 8 | 2620.44 | 327.56 | 1.31 | 0.247 |
| Series within Element | 14 | 6609.16 | 472.08 | 1.89 | 0.036 |
| Error | 105 | 26270.46 | 250.19 |  |  |
| Total | 133 | 53987.24 |  |  |  |

**Figure 2.**
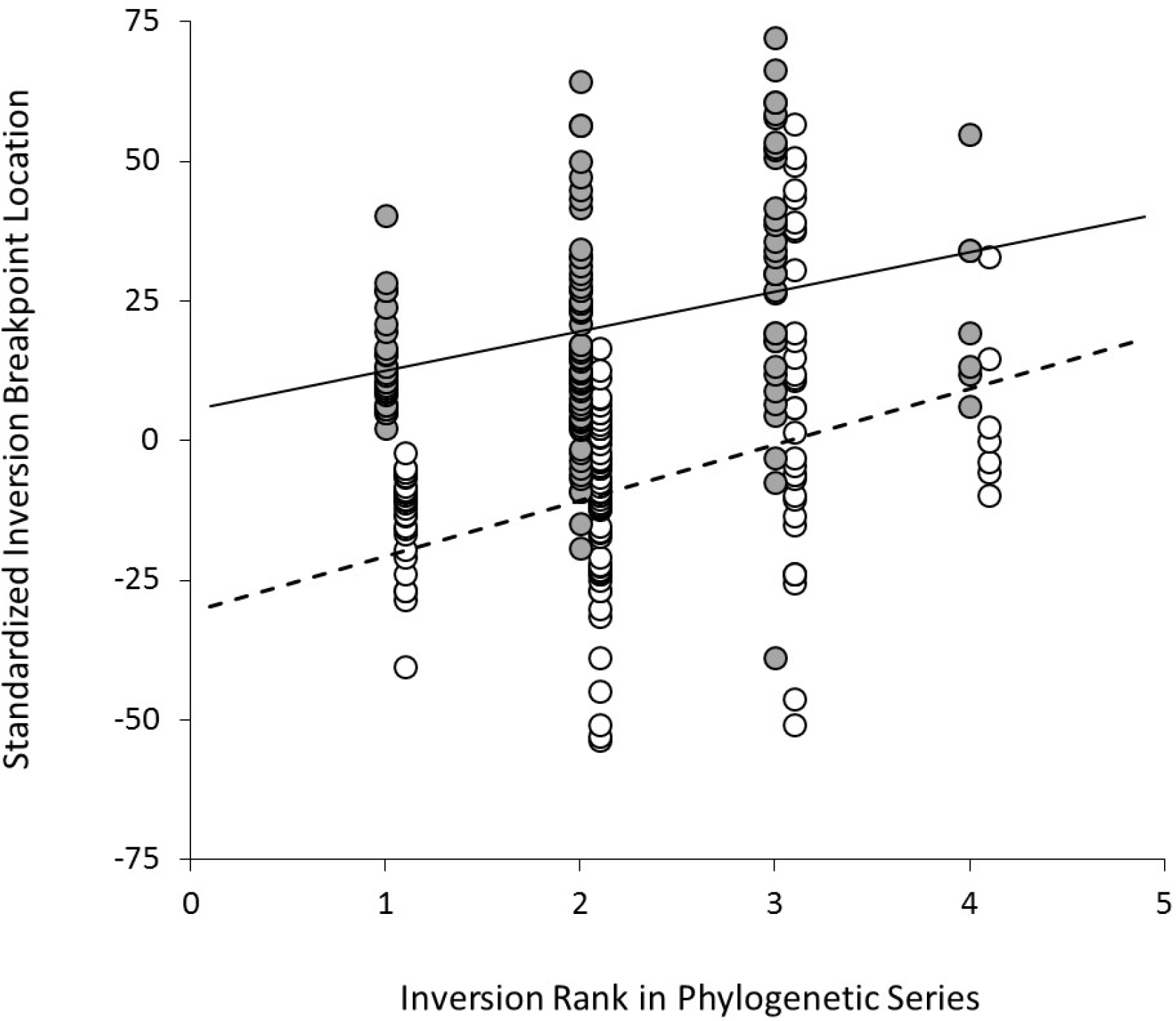
Standardized inversion breakpoint location regressed on phylogenetic rank. Movement in the positive direction on the standardized scale is a movement toward the telomere (a distal shift). Closed circles are the distal breakpoints, open circles are proximal breakpoints (displaced by +0.1 units on the x-axis for ease of visualization). Statistically significant linear regression is depicted by solid line for distal breakpoints and dotted line for proximal breakpoints (table 2 and 3).

**Table 3.** ANOVA table for regression of distal inversion breakpoint location on phylogenetic rank. Data corresponds to the shaded symbols in figure 2

| Source of Variation | *df* | *SS* | *MS* | *F ratio* | *p* |
| --- | --- | --- | --- | --- | --- |
| Regression on Rank | 1 | 2574.71 | 2574.71 | 8.86 | 0.004 |
| Species | 5 | 992.29 | 198.46 | 0.68 | 0.637 |
| Element within Species | 8 | 3008.67 | 376.08 | 1.29 | 0.254 |
| Series within Element | 14 | 5927.14 | 423.37 | 1.46 | 0.140 |
| Error | 105 | 30499.63 | 290.47 |  |  |
| Total | 133 | 49107.05 |  |  |  |

The telomeric progression was stronger for the proximal breakpoints than the distal breakpoints (figure 2), suggesting a reduction in size for inversions of high rank (figure 3). However, the slopes of these two regressions do not differ with statistical significance (*F_1,4_=0.37, p=0.496*) (table 4), and the apparent trend towards size reduction was not statistically significant (*β=-2.93, t=-1.25, df=1 p=0.22*) (table 5). Empirically, the phylogenetic series from just six species provides strong evidence for distal shift but insufficient data to demonstrate the trend toward size reduction with statistical significance.

**Table 4.** ANOVA table testing equality of slopes from regression of proximal and distal inversion breakpoints against phylogenetic rank. The weighted average deviation from regression 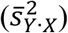 was calculated as described in Sokal and Rohlf (1995) using summary statistics from tables 2 and 3

| Source of Variation | *df* | *SS* | *MS* | *F ratio* | *p* |
| --- | --- | --- | --- | --- | --- |
| Variation among regressions | 1 | 398.02 | 398.02 | 0.37 | 0.496 |
| Average variation within regressions | 4 | 4328.72 | 1082.18 |  |  |

**Figure 3.**
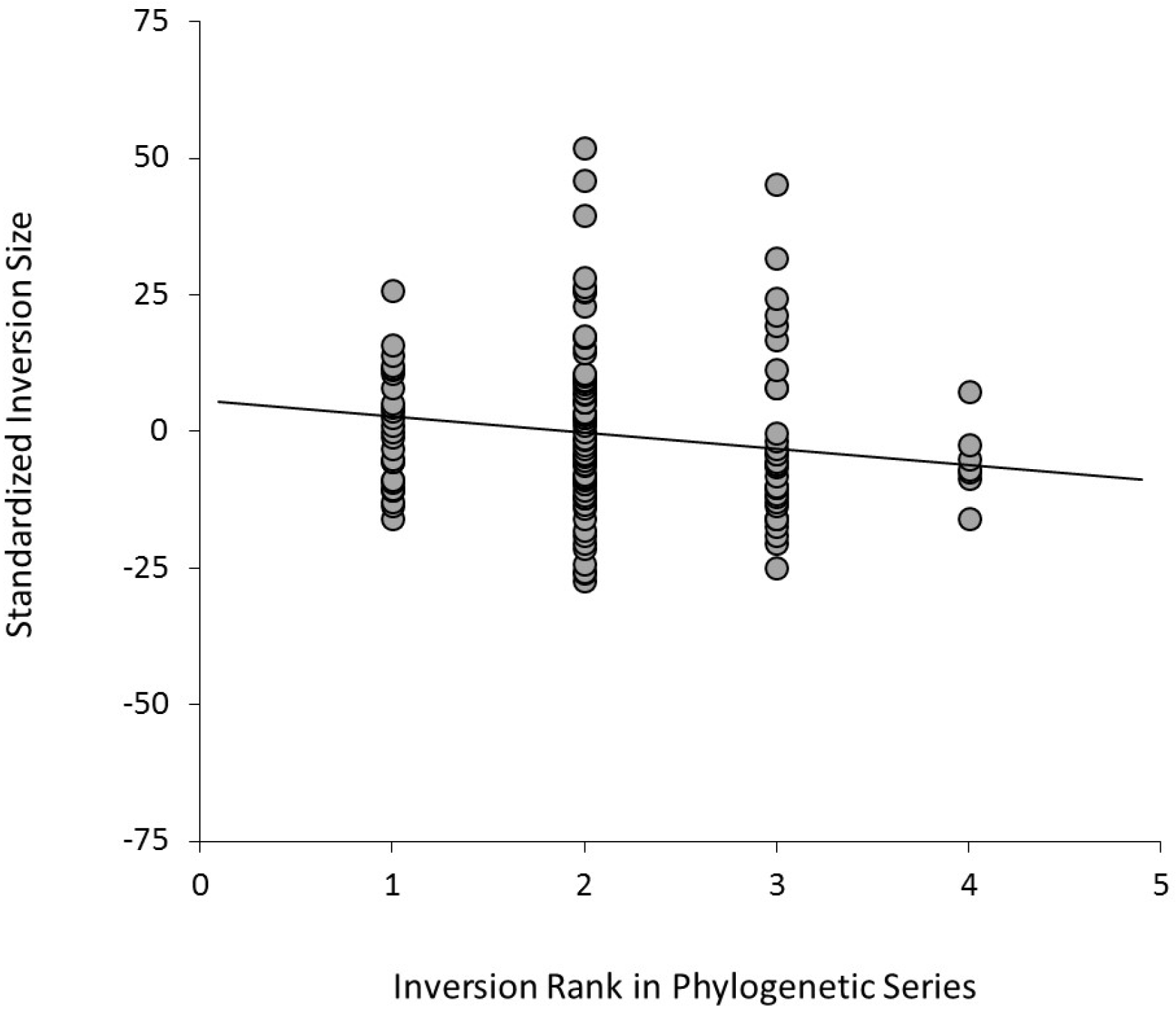
Progressive trend towards inversion size reduction in phylogenetic series. The line of best fit for inversion size regressed on phylogenetic rank is depicted by solid line (table 5). Regression coefficient of this line does not differ from zero with statistical significance.

**Table 5.** ANOVA table for regression of inversion size on phylogenetic rank. Data corresponds to figure 3.

| Source of Variation | *df* | *SS* | *MS* | *F ratio* | *p* |
| --- | --- | --- | --- | --- | --- |
| Regression on Rank | 1 | 441.62 | 441.62 | 1.55 | 0.216 |
| Species | 5 | 16.63 | 3.33 | 0.01 | > 0.999 |
| Element within Species | 8 | 92.43 | 11.55 | 0.04 | > 0.999 |
| Series within Element | 14 | 94.32 | 6.74 | 0.02 | > 0.999 |
| Error | 105 | 29886.67 | 284.63 |  |  |
| Total | 133 | 30355.41 |  |  |  |

## DISCUSSION

### Novitski’s Distal Shift

The telomeric progression of overlapping inversions for serially inverted chromosomes was previously known only as qualitative pattern from a single Muller element. The distal shift was confirmed here by a quantitative analysis of 28 phylogenetic series of 134 paracentric inversions from six *Drosophila* species of the *obscura* group. Each phylogenetic series arose independently in every species and represents the direct ancestor-descendent relationships in paracentric inversion evolution. Therefore, each of the 28 phylogenetic series is a unique, independent realization of a distinct directional pattern in chromosome evolution. To what degree the distal shift represents a general rule of inversion evolution, as opposed to a chromosomal anomaly of the *D. obscura* group, is the subject of a forthcoming publication.

One feature of the *obscura* group sex chromosomes proves an exception to this rule. “Sex ratio” (SR) chromosomes are coadapted gene complexes that cause strong unequal transmission of X and Y chromosomes and are found in all six species (Jaenike, 2001). SR chromosomes in all six species carry inverted gene arrangements, but these inversions tend to be non-overlapping and rarely form phylogenetic series. When SR chromosomes do form phylogenetic series (Muller element A of *D. subobscura,* element D of *D. athabasca*), they tend to be smaller series and do not exhibit the distal shift or size reduction. It is likely that the very strong selection on sex ratios and recombination suppression associated with the strong transmission bias of SR chromosomes overwhelms the statistical signal produced the relatively weak force that drives the distal shift.

If the distal shift observed for autosomes in these species is just an extreme illustration of some underlying principle common to all paracentric inversion evolution, then a cytogenetic mechanism is required for this pattern. Novitski himself presented a biased mutational model after observing the non-uniform distribution of inversion breakpoints in the phylogenetic series of *D. pseudoobscura* (Bernstein and Goldschmidt, 1961; Novitski, 1946; Novitski, 1961). This ingenious model invokes a bias of spontaneous chromosome breakage in inversion heterozygotes and remains a viable explanation awaiting experimental examination. However, as Novitski noted, this model is limited to explaining the clumped distribution of inversion breakpoints and alone is insufficient to explain the distal shift, size reduction, or variability of inversion abundance (Novitski, 1946). Below I present an alternative mechanism with well-validated assumptions that addresses all these aspects of inversion polymorphism, and does not invoke the logistically untestable mutational bias assumption.

### Meiotic Drive Mechanism

In *Drosophila*, recombination between different gene arrangements is effectively suppressed because crossing over produces acentric and dicentric meiotic products that for mechanical reasons are relegated to the polar body nuclei and never included in the functional egg (Hinton and Lucchesi, 1960; Sturtevant and Beadle, 1936). However, as illustrated in figure 4A, heterozygotes for overlapping inversions have a homosequential region, where if crossing over were to occur, all four meiotic products would be monocentric and could in principle be included in the functional egg (figure 4B). Because of the figure eight pairing pattern (figure 4C), crossing over in meiosis I generates large deletions and duplications resulting in dyad asymmetry (figure 4B) (Sturtevant and Beadle, 1936). In *Drosophila* females, it is a thoroughly established fact that the shorter chromatid of an asymmetric dyad in meiosis II has a higher probability of being included in the functional egg (Lindsley and Sandler, 1965; Novitski and Sandler, 1956; Zimmering, 1955). The phenomenon of unequal recovery from asymmetric dyads is known as nonrandom disjunction and is a well-known form of female meiotic drive (reviewed in Novitski, 1951; Novitski, 1967).

**Figure 4.**
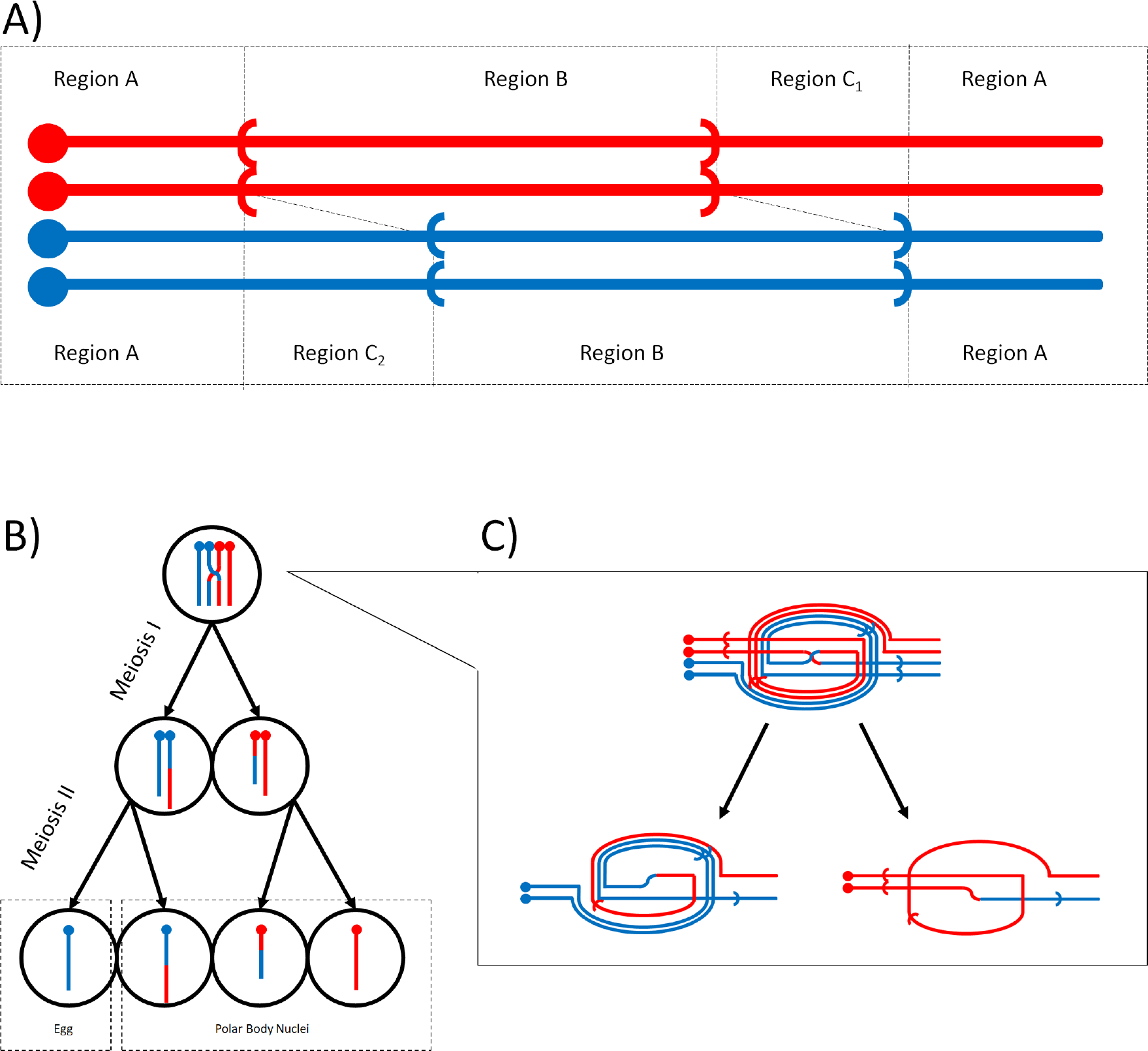
Schematic for nonrandom disjunction of overlapping inversion. A) Four strand bundle diagram divided into homosequential regions external to inversions (Region A), regions inverted relative to one another (Region C_1_ and C_2_), and the homosequential region internal to inversions (Region B). B) Progression of asymmetric products from a crossover in Region B through meiosis I and II, migration of chromatids to the egg pole is probabilistic but always favors the transmission of distal inversion (blue non-recombinant chromatid). C) Figure eight pairing of overlapping inversions and the resulting asymmetric dyads resulting from crossing over in Region B, illustrating all four possible meiotic products and their relative sizes.

For any overlapping paracentric inversions that are two steps apart in a phylogenetic series, a homosequential region exists where crossing over produces asymmetric dyads (Sturtevant and Beadle, 1936). Furthermore, for any such inversions, a single crossover event will create dyads pairing the distal inversion with the duplications and the proximal inversion with the deletions. The resulting asymmetric dyads will favor the inclusion of the chromatids carrying the distal inversion and the deletions in the functional egg. Because large chromosomal deletions and duplications generally form inviable zygotes, the final result is an over-representation of distal inversions in the viable progeny of females heterozygous for overlapping inversions.

This meiotic drive mechanism applies equally to included inversions (supplemental figure 2), where the asymmetric dyads generated from crossing over consist of a proximal duplication and distal deletion (or vice versa). The mechanism operates for inversions in repulsion phase (as illustrated in figure 4) or coupling phase as observed for serially inverted chromosomes (supplemental figure 3 and 4). In the latter case, the serially inverted chromosome will drive against the standard arrangement only if the second inversion is distal to the first. Therefore, single crossover events in homosequential regions of any two inversions (overlapping or included, in repulsion or coupling phase) will unequivocally create a bias favoring the inclusion of distal inversions in the functional egg. I propose that it is this intrinsic bias in female meiosis that generates the distal shift in a phylogenetic series of serially inverted chromosomes. This hypothesis also explains the weak tendency towards size reduction as a by-product of favoring evermore distal inversions in subtelomeric regions where large inversions are precluded by the position of the telomere.

### Challenges of Drive Theory

This meiotic drive theory of Novitski’s distal shift presents several difficulties from both transmission genetic and population genetic perspectives. Although crossing over in homosequential regions of overlapping inversions has been directly observed, experimental investigation has been limited to X chromosomes and requires the use of compound chromosomes or translocation stocks to recover recombinant products (Grell, 1962; Novitski and Braver, 1954; Sturtevant and Beadle, 1936). Crossing over in shared inverted regions has not yet been demonstrated for inversions of autosomes segregating in natural populations. Formal genetic analysis for the common inversions of Muller element E in *D. melanogaster* and Muller element C in *D. pseudoobscura* could not detect nonrandom disjunction (Meisel and Schaeffer, 2007, Koury unpublished). However, using realistic parameters for crossing over and nonrandom disjunction, expected transmission ratios do not exceed *k* = 0.513 (Koury unpublished), a deviation that is on the same order of magnitude of viability effects of phenotypic markers used and well within the measurement error of both experiments. Nonetheless, it is not uncommon for effects below the threshold of experimental detection (*e.g.* codon bias) to have major evolutionary significance. Furthermore, there are experimental refinements possible with *D. melanogaster* model system currently being pursued to enhance the ability to experimentally detect drive in this scenario.

More difficulty is encountered when considering the population genetics of the distal shift. The relative rarity of inversion mutations, overlapping inversion heterozygotes, and crossing over in homosequential regions (each a precondition for the next) paired with the relatively weak strength of drive and the underdominance of overlapping inversion heterozygotes (due to dominant lethal deletion and duplications), suggests a small role for nonrandom disjunction in paracentric inversion evolution. In considering the extension to serially inverted chromosomes, the later gene rearrangements can only drive at the expense of the earlier steps. So this force, while of plausible importance in the first, second, or third steps of a phylogenetic series, quickly becomes vanishingly small in later steps. These population genetic questions require rigorous quantitative analysis beyond the scope of this paper and are the subject of a forthcoming publication.

Interestingly, the challenges outlined in this section generate several predictions which are consistent with the *obscura* group data. First, the relatively weak female meiotic drive for overlapping inversions (maximum *k* = 0.513) does not generate a distal shift for SR chromosomes where stronger forces are expected to prevail. Second, for the most extensive and best resolved phylogenetic series (Muller element C of *D. athabasca, D. pseudoobscura*, and *D. subobscura*), the ancestral arrangements and early steps tend to be rare or absent. Finally, for autosomes no series greater than four inversion steps was observed (supplement table 1) and the distal shift is relatively weak for the few inversions of rank four (figure 3).

### Patterns in Inversion Evolution

It is very encouraging to note that the proposed meiotic drive mechanism bears on several other patterns in inversion polymorphism. To justify experimental investigation and population genetic simulations for this scenario I enumerate some of these observations. The patterns of paracentric inversion variability mentioned in the introduction suggest this form of structural heterozygosity has autocatalytic properties (Bernstein and Goldschmidt, 1961; Novitski, 1961). The meiotic drive mechanism predicts serial inverted chromosomes, especially distally placed second inversions, have intrinsic advantages in invading a population already segregating for chromosomal rearrangements in the same genomic region. And although this advantage does not hold for advanced stages in the phylogenetic series, the approach to complete recombination suppression by favoring inversions in later stages has the second order effect of reducing genetic load due to this particular form of meiotic drive (Crow and Kimura, 1970).

In considering just three consecutive steps of a phylogenetic series (ancestral, intermediate, and derived), Wallace observed that the intermediate arrangement is often absent in a given population (Wallace, 1953). Wallace’s “Rule of Triads” is immediately comprehensible on the view that the intermediate and derived arrangements have similar gene contents and fitness; however, the derived arrangement has the added benefit of driving against the ancestral state and thus outcompetes the intermediate arrangement. Along these same lines, the commonly observed local extirpation of arrangements of low rank in a long phylogenetic series may be related to being commonly driven against.

Finally, although Novitski’s (1961) biased mutational model explains the clustered pattern of inversion breakpoints generating extensive inversion overlap, the meiotic drive theory offers an equally viable alternative. The drive mechanism predicts, even with uniform distribution of spontaneous inversion breakpoints, that the inversions with greatest overlap and thus greatest opportunity for nonrandom disjunction, would invade natural populations. The clumped distribution of observed breakpoints would therefore be the result of biases during the establishment phase of inversion not from any spontaneous mutational bias.

Rates of spontaneous chromosomal inversion, and any biases thereof, are outside the scope of reasonable experimental investigation (*cf*. Yamaguchi and Mukai, 1974 for an unreasonable attempt). As a consequence it is unclear how to practically differentiate alternative theories of chromosome evolution based solely on patterns of natural inversion polymorphism. The meiotic drive theory of paracentric inversion evolution introduced here has the potential to explain with a single mechanism a number of different chromosome patterns that were previously thought to be unrelated. Furthermore, the meiotic drive theory is based on a cytogenetic mechanism that is amenable to direct experimentation, thereby conferring a high degree of testability to this model of chromosome evolution.

### Conclusion

In a phylogenetic series of overlapping inversions in the *Drosophila* species of the *obscura* group derived arrangements tend to have distally shifted breakpoints resulting in smaller inversions. The distal shift, viewed *in extremis* for the hypervariable Muller element C of *D. pseudoobscura* and *D. athabsaca*, likely reflects a fundamental mechanism of paracentric inversion evolution, while the size reduction is simply a byproduct of the distal shift in subtelomeric regions. Nonrandom disjunction of overlapping inversions was demonstrated to always favor transmission of distal inversions and is hypothesized here to favor the evolution of serially inverted chromosomes. Therefore, far from being selectively beneficial, inversion polymorphism, serially inverted chromosomes, and the associated distal shift result from intrinsic biases in meiosis and generate a substantial genetic load. This novel hypothesis requires further investigation along both experimental and theoretical lines. The meiotic drive mechanism proposed should be of considerable interest as it can explain Novitski’s distal shift as well as several related patterns of breakpoint distribution and paracentric inversion evolution.

## Supporting information

