## Supplementary material for "Novitski’s Distal shift in Paracentric Inversion Evolution"

Supplemental Figure 1.  Illustration of all full phylogenetic series used in this study.

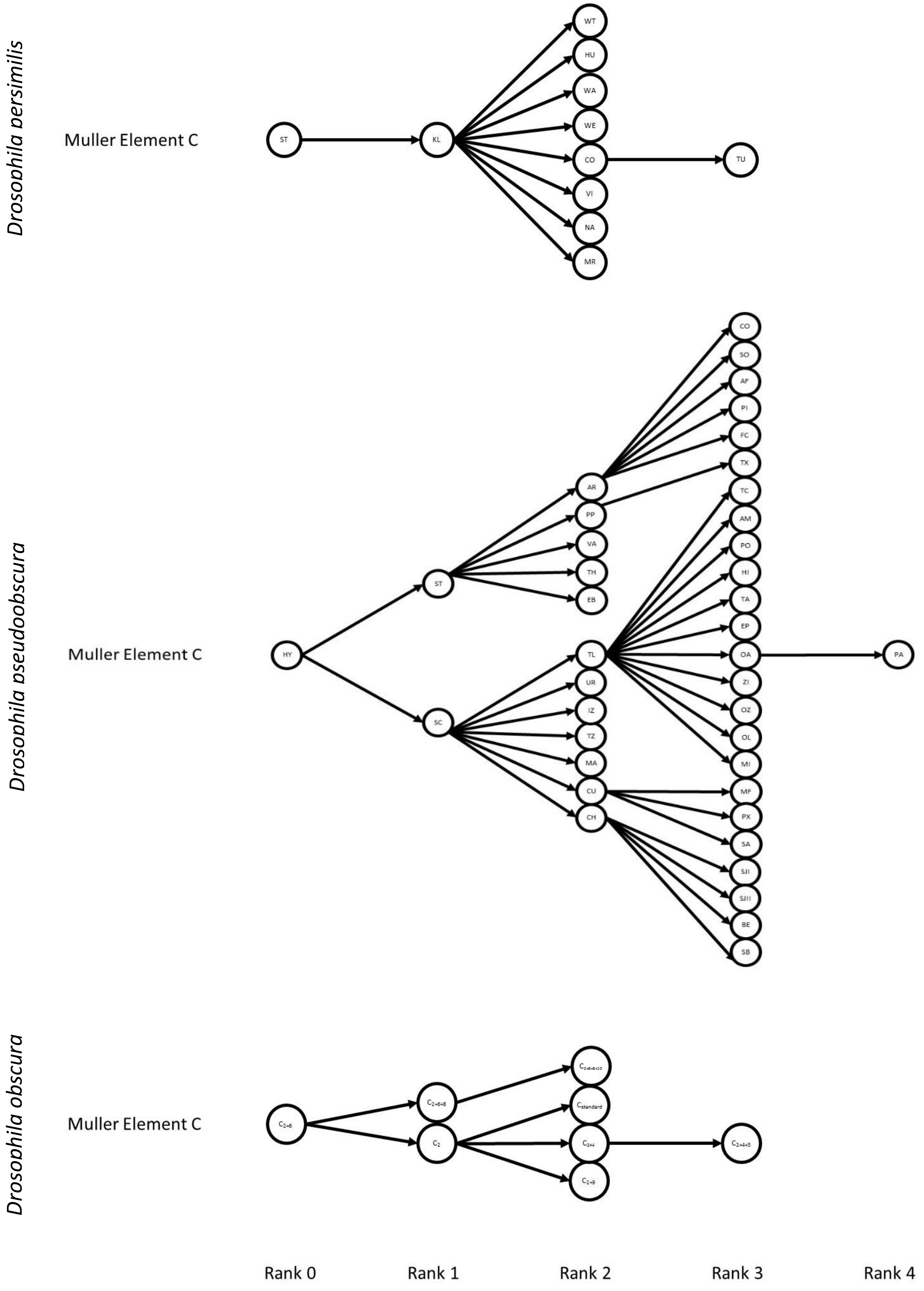

Supplemental Figure 1 continued.  Illustration of all full phylogenetic series used in this study.

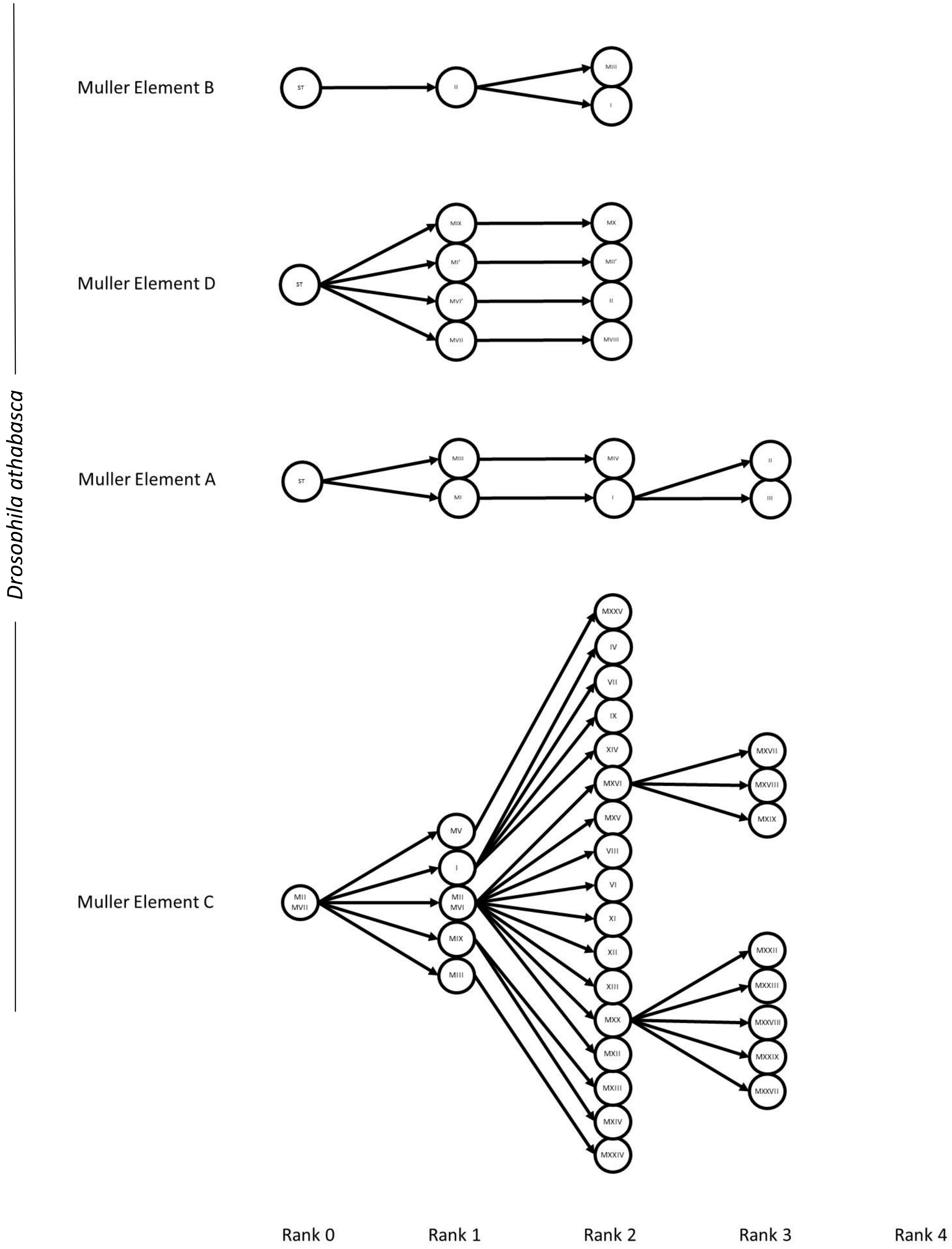

Supplemental Figure 1 continued. Illustration of all full phylogenetic series used in this study.

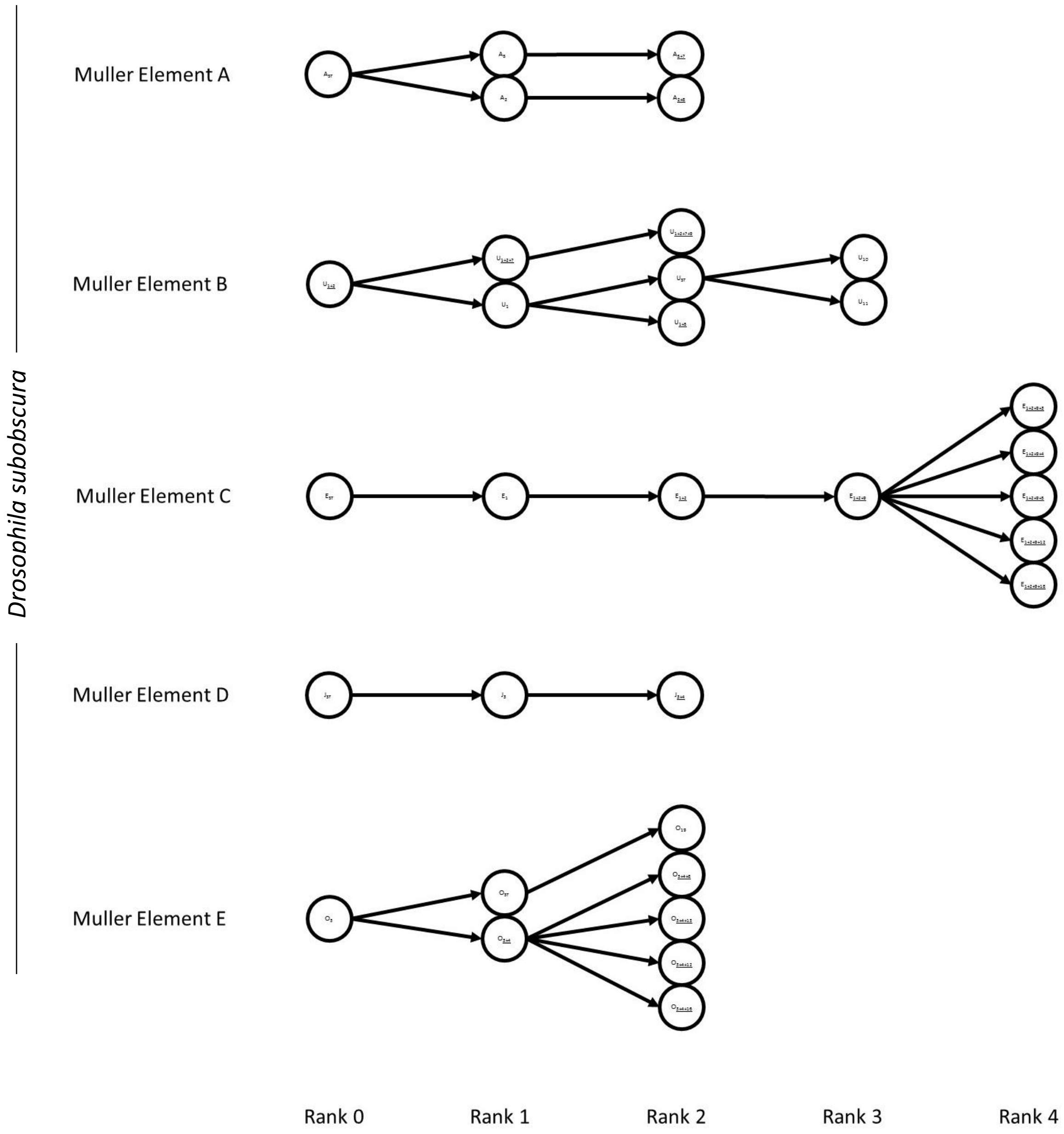

Supplemental Figure 1 continued.  Illustration of all full phylogenetic series used in this study.

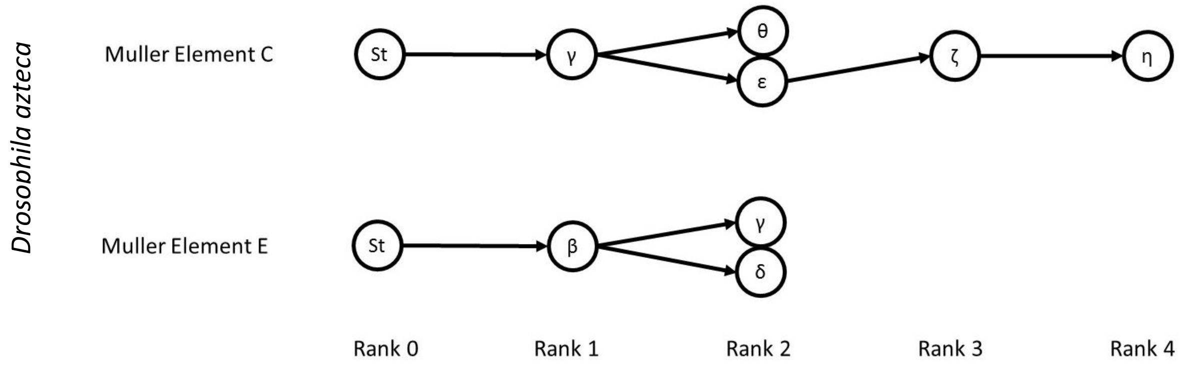


Supplemental Figure 2. Pairing diagrams for included inversions in repulsion phase.

Meiotic products for crossover located in Region B, size asymmetry illustrated is proportional.

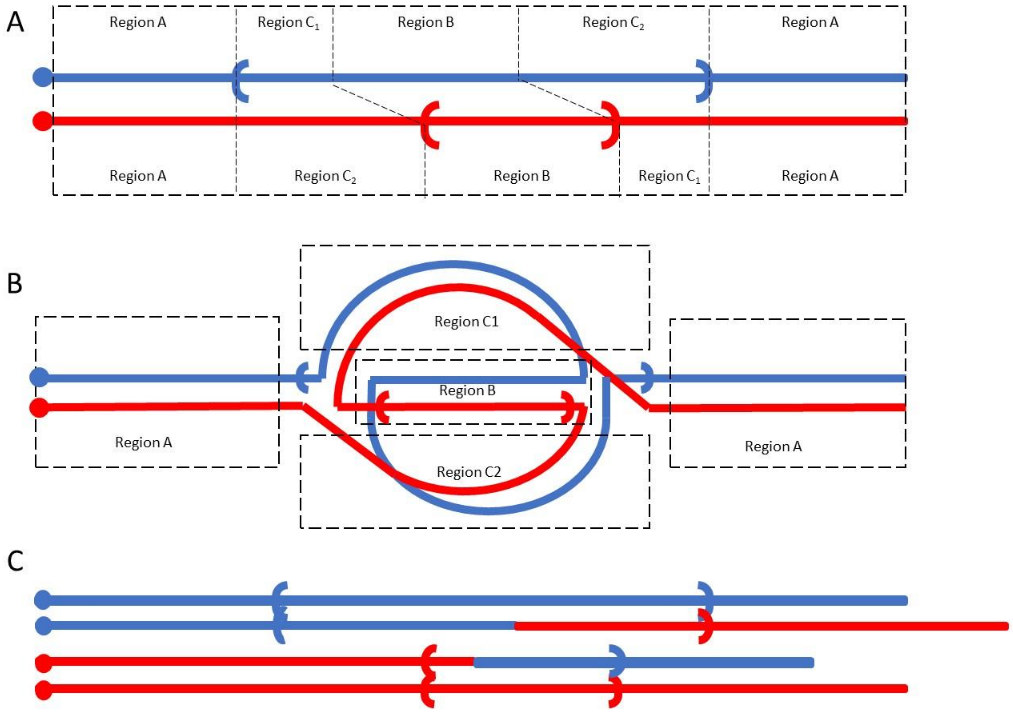

Supplemental Figure 3. Pairing diagrams for overlapping inversions in coupling phase.

Meiotic products for crossover located in Region B, size asymmetry illustrated is proportional.

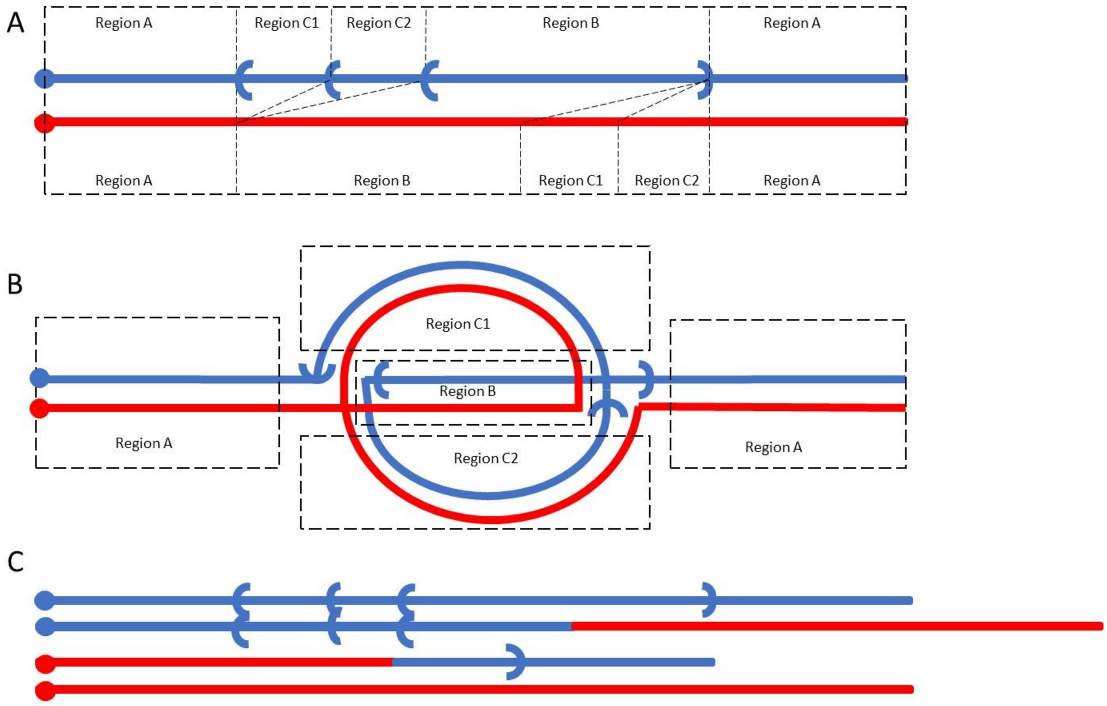


Supplemental Figure 4. Pairing diagrams for included inversions in coupling phase.

Meiotic products for crossover located in Region B, size asymmetry illustrated is proportional.

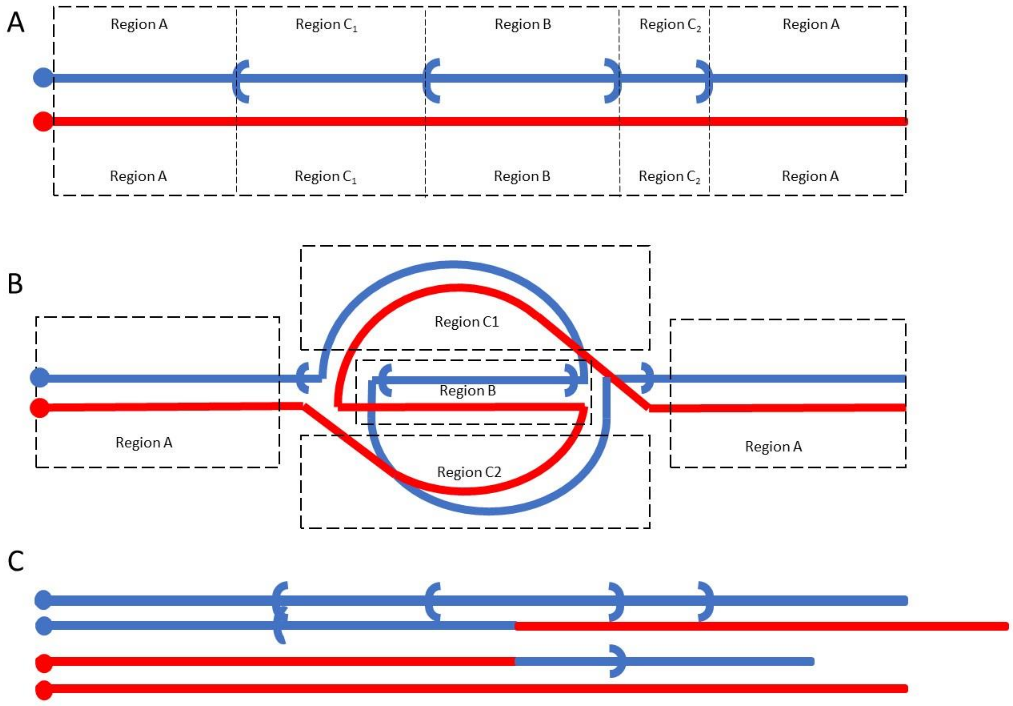
